# Probing circuit mechanisms of feature selectivity in mouse visual cortex through synaptic-resolution connectomics

**DOI:** 10.1101/2024.11.18.624135

**Authors:** Victor Buendía, Jacopo Biggiogera, Alessandro Sanzeni

## Abstract

Feature selectivity, the ability of neurons to respond preferentially to specific stimuli, is a defining property of cortical computation. Competing theories attribute selectivity to structured, tuning-dependent “like-to-like” feedforward or recurrent connections, whereas others propose that it can emerge without specific structure in randomly connected, inhibition-dominated networks. The relative contribution of these mechanisms remains unclear. Here, we investigate the circuit basis of feature selectivity in mouse visual cortex, focusing on how orientation selectivity arises in layer 2/3 neurons driven by input from layer 4. We developed a data-driven modeling framework integrating network modeling with functional imaging and synaptic-resolution connectomics from the MICrONS dataset. Our analyses show that randomness in connectivity is the dominant source of selectivity, while structured “like-to-like” feedforward and recurrent connections play comparable secondary roles in amplifying it. These findings refine classical theories of cortical selectivity and demonstrate how connectome-constrained modeling can reveal the circuit principles underlying cortical computation.

---

Neural responses exhibit selectivity for specific stimulus features across sensory areas, such as edge orientation in primary visual cortex (V1) [1], sound frequency in auditory cortex [2], and odor identity in piriform cortex [3]. Beyond sensory coding, selectivity is a hallmark of diverse brain functions, including movement planning, spatial navigation, and social cognition [4–8]. The form of selectivity often evolves across successive processing stages. In the primate visual system, for instance, it progresses from orientation tuning in V1 to corners in secondary areas and to complex features such as objects and faces in inferotemporal cortex [9–12]. These hierarchical transformations are thought to support increasingly abstract representations that enable advanced visual computations, including invariant object recognition [13–15].

The very presence of stimulus selectivity within highly recurrent cortical networks presents a theoretical puzzle. In such networks, where neurons integrate a large number of stimulus-dependent inputs, heterogeneity in connectivity can dilute information and lead to a loss of selectivity (see [16, 17] and derivation in Supp. Sec. I). This raises a fundamental question: how does selectivity emerge in the cortex? Three influential classes of models have been proposed to address this question. In *feedforward models*, selectivity is inherited through structured connections that pool inputs from neurons with specific tuning [1, 18]. In *recurrent models*, weakly selective inputs are amplified by recurrent interactions in which neurons with similar response properties are more strongly interconnected [19–21]. Finally, in *balanced models*, networks of randomly connected neurons acquire selectivity through strong recurrent inhibition that cancels the dominant non-selective input, allowing weaker stimulus-dependent components to drive neural responses [16, 17]. While both feedforward and recurrent models emphasize structured wiring, balanced models suggest that selectivity can also emerge in randomly connected networks.

Experimental studies in the visual cortex have reported structured connectivity, whereby neurons with similar orientation preferences are more likely to be connected, a pattern commonly referred to as “like-to-like” [22–27]. Such findings support models in which tuning-dependent connectivity generates selectivity through structured feedforward input, recurrent amplification among similarly tuned neurons, or both. In mouse V1 layer 2/3, presynaptic partners in layers 2/3 and 4 exhibit similar tuning properties [27, 28], suggesting that both pathways contribute to orientation selectivity, although their respective roles and relative contributions to postsynaptic tuning remain poorly constrained. Moreover, while experiments describe enhanced connectivity among similarly tuned neurons, they also revealed that a substantial fraction of connections link neurons with dissimilar tuning, reflecting the random component of cortical wiring. Additional variability arises from the heterogeneous distribution of synaptic strengths [29, 30] and neuronal response properties. Modeling studies have shown that such heterogeneity alone can give rise to selectivity [16, 17], consistent with experimental data [31], and can even produce emergent “like-to-like” connectivity patterns [17]. It therefore remains unclear to what extent structured connectivity is necessary for generating selectivity and how feedforward and recurrent pathways interact with random connectivity to shape this process.

A promising way to test competing models of feature selectivity is to directly constrain circuit models with connectomic data. Recent advances in electron microscopy (EM) have enabled the reconstruction of complete wiring diagrams of neural tissue at synaptic resolution [32–37]. This progress has opened new opportunities for model construction, with successful applications in non-cortical systems such as the retina, olfactory bulb, and insect brain [38–46]. Extending such approaches to cortex has only recently become feasible through large-scale EM reconstructions. Yet current connectomic datasets remain limited by missing physiological data and incomplete reconstructions [47, 48], underscoring the need for methods that integrate structural data with functional and theoretical modeling.

In this study, we investigate the circuit mechanisms underlying feature selectivity in mouse V1, focusing on how orientation selectivity arises in layer 2/3 (L2/3) neurons in response to input from their primary driver, layer 4 (L4). To this end, we developed a datadriven modeling framework that leverages the MICrONS dataset [36], which combines functional imaging during visual stimulation with a dense, synapse-resolution connectome from the same cortical volume. This dataset provides the most comprehensive co-registered structural and functional reconstruction of mouse visual cortex to date, offering an unprecedented basis for mechanistic models of selectivity.

Our work addresses two central questions: how feedforward input from L4 and recurrent input within L2/3 contribute to selectivity, and how structured connectivity and random variability shape this process. By building models directly constrained by structural and functional data, we quantitatively reproduce the selectivity observed experimentally. We find that randomness in connectivity is the dominant driver of selectivity, while structure in feedforward and recurrent connections from L4 and L2/3 plays comparable secondary roles in amplifying it. More broadly, our framework demonstrates how connectomic reconstructions can uncover the circuit mechanisms that underlie cortical computation.

## RESULTS

### Constraining models of feature selectivity with connectomes: challenges and approaches

We characterized neuronal selectivity in L2/3 and L4 of mouse V1 using the functional component of the MICrONS dataset [36], which includes responses from thousands of excitatory neurons (4620 in L2/3 and 3139 in L4) recorded during presentations of parametric and naturalistic visual stimuli (see Methods). A large fraction of neurons in both layers were orientation selective (78% in L2/3 and 67% in L4; example tuning curves in Fig. 1a), consistent with previous reports [49, 50]. Preferred orientations were unevenly distributed, with peaks near 0 and *π/*2 (Fig. 1b), in line with large-scale recordings [51, 52] and potentially reflecting adaptation to natural image statistics [53]. Average tuning curves were similar across layers (Fig. 1c), but neurons exhibited broad variability in selectivity, quantified by the circular variance (Fig. 1d). Non-selective neurons showed diverse responses, comparable to those of selective neurons at non-preferred orientations.

**FIG. 1.**
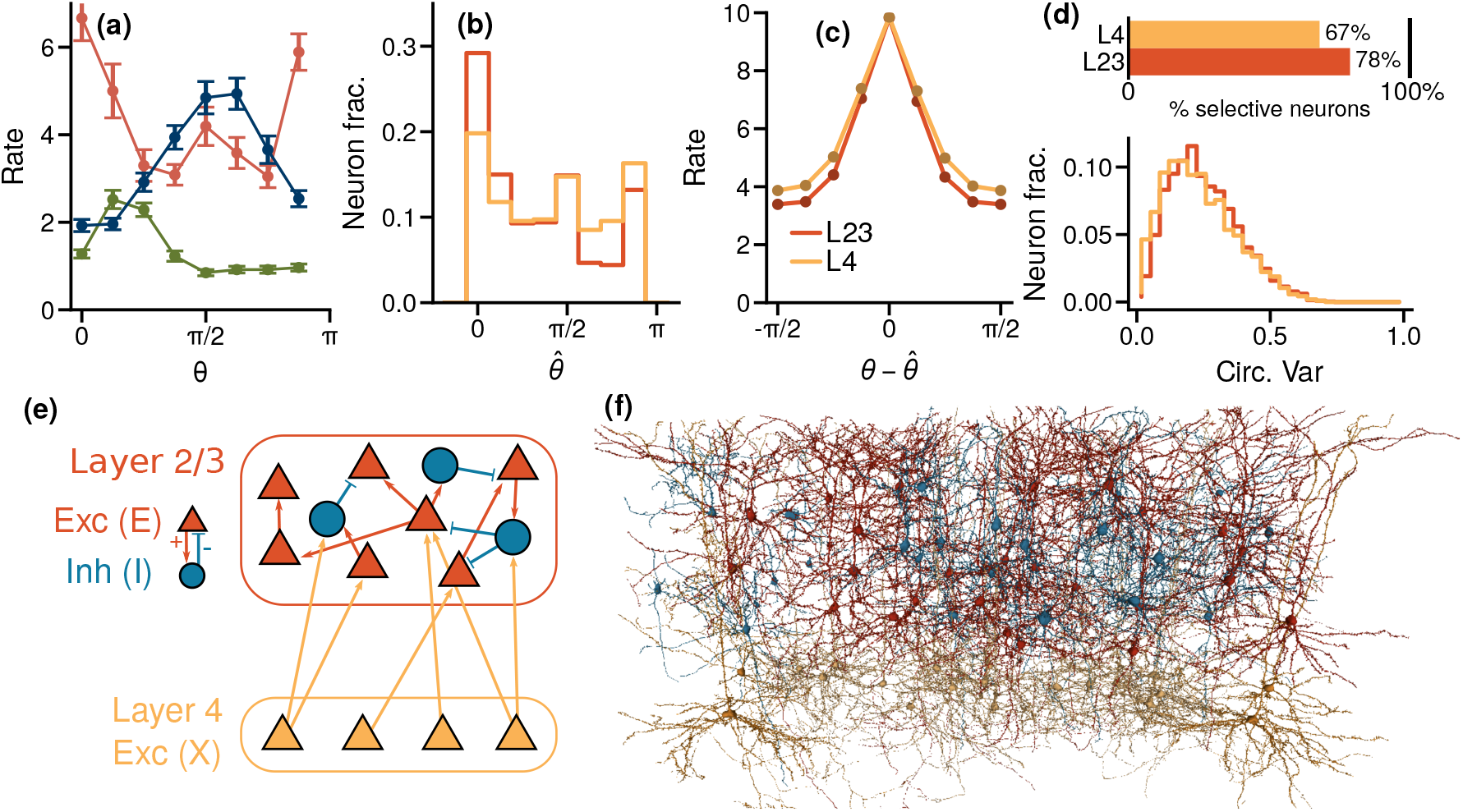
Investigating the emergence of orientation selectivity using connectome-constrained network modeling. Orientation tuning curves of example excitatory neurons in layers 2/3 (L2/3) and 4 (L4) of mouse V1 from the functional component of the MICrONS dataset [36]. **(b)** Distributions of preferred orientations for L2/3 and L4 neurons. **(c)** Population-averaged tuning curves for orientation-selective neurons in each layer. (**d**) Selectivity metrics: fraction of tuned neurons (top) and distribution of circular variance (bottom; see Methods for definition, values closer to 0 and 1 indicate weak and strong tuning, respectively). **(e)** Schematic of the recurrent network model (Eq. (1)), with L4 excitatory input (X) driving L2/3 excitatory (E) and inhibitory (I) populations. **(f)** Electron microscopy reconstructions of L2/3 and L4 neurons from the MICrONS structural dataset [36]. Functional recordings (**a–d**) and connectomic data (**f**) were combined to build network models (**e**) that reveal how orientation selectivity in L2/3 emerges from L4 input and local recurrent interactions.

We modeled neural responses in L2/3 to L4 inputs using networks of recurrently connected excitatory (*E*) and inhibitory (*I*) neurons, representing L2/3 populations, driven by an excitatory population (*X*) modeling L4 excitatory neurons (Fig. 1e,f). In the model, the orientation *θ* of the stimulus determines the firing rates 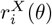 of the *X* neurons, while the rates 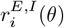 are shaped by recurrent dynamics given by

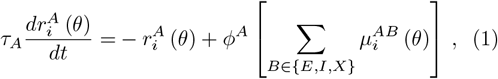

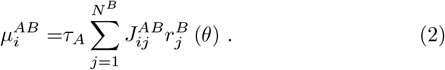

Here, φ^*A*^(·) and *τ*_*A*_ are the single-neurons input-output function (firing rate vs. input current) and the time constant of population *A* ∈ {*E, I* }, respectively. The inputoutput response follows a Ricciardi nonlinearity [54] (see also Methods). 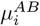 is the current into neuron *i* of population *A* ∈ {*E, I*} from population *B* ∈ {*E, I, X* }, calculated as the sum of individual inputs 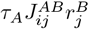 from the *j*-th neuron in the presynaptic population, along with the synaptic efficacy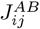.

If all components of Eq. (1) were experimentally accessible, the model could be directly tested to dissect the circuit mechanisms underlying feature selectivity. The MICrONS dataset [36] provides valuable constraints but does not fully specify the model. In addition to its functional recordings, it includes a structural reconstruction with 22,647 excitatory neurons (9,487 in L4 and 13,160 in L2/3) and 1,299 inhibitory neurons in L2/3, where synapse size serves as a proxy for synaptic efficacy 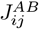 [55]. Key elements remain unconstrained: synapse sizes determine strengths only up to an unknown scaling factor [55], which can strongly affect network dynamics [56, 57]. Neuron-specific properties (e.g., φ^*A*^, *τ*_*A*_) are not available, and only a subset of excitatory neurons are matched across modalities (35% in L2/3, 33% in L4), with inhibitory responses entirely absent. Automated reconstructions also introduce connectivity errors [48]; although manual proofreading reduces these, it has been applied to only 972 neurons, including 453 in L2/3.

The limited constraints on Eq. (1) underscore the need for methods that maximize the utility of the MICrONS dataset [36] while addressing its gaps. To this end, we developed two complementary strategies to investigate the mechanisms of selectivity. First, we analyzed excitatory synaptic currents, which are insensitive to unmeasured factors such as single-neuron input-output properties. Second, we constrained recurrent network models with the dataset, inferred missing components using Bayesian approaches, and performed simulations and perturbation analyses to probe circuit mechanisms.

### Feedforward and recurrent inputs contribute similarly to orientation selectivity

Feedforward and recurrent models of cortical computation make distinct predictions about the origin of orientation-tuned input. Feedforward models posit that a large, sharply tuned excitatory drive arises from L4 and drives L2/3 responses [1, 18], whereas recurrent models propose that tuning is generated within L2/3 through amplification of feedforward input by local interactions [19–21]. To test these alternatives, we derived from Eq. (1) its stationary form, which links each neuron’s firing rate to the sum of its synaptic inputs:

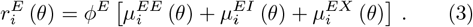

Although the input–output function φ^*E*^ is not directly constrained, it is monotonic [58], implying that the orientation dependence of firing is determined by the total synaptic input. Therefore, analyzing synaptic currents enables us to assess feedforward and recurrent contributions without specifying the exact form of φ^*E*^.

A direct consequence of Eq. (3) is that a neuron’s preferred orientation should correspond to the orientation at which its total input is maximal. To test this, we estimated the total excitatory drive to each orientation-selective L2/3 neuron by summing the product of each presynaptic partner’s tuning curve and synaptic size. This computation combined structural and functional data from the MICrONS dataset [36], using synapse size as a proxy for excitatory strength. Inhibitory currents could not be estimated directly because inhibitory recordings are unavailable [36], but previous studies show that inhibitory neurons are less selective than excitatory ones and that their tuning is typically broader [27, 49, 59, 60]. Their omission is therefore expected to have only a minor effect on the orientation dependence of postsynaptic responses, an issue explicitly examined in subsequent network analyses that included inhibitory dynamics.

Figure 2a-c shows the orientation dependence of firing rate and excitatory input computed for an example neuron using Eq. (3). As expected, the estimated input peaked at the neuron’s measured preferred orientation. This pattern held across the population: average excitatory input peaked near the postsynaptic preference (Fig. 2d,e), and the orientation inferred from total input matched the measured value on average (Fig. 2f,g). Alignment was imperfect, with many neurons deviating from the predicted relationship between input and firing preference. This likely reflects the small number of synaptic inputs available per neuron, resulting from automated segmentation errors and the restriction of the analysis to presynaptic neurons with manually proofread axons [61] and functional matches. On average, L2/3 excitatory neurons received 6.71 *±* 0.09 identified inputs, only a small fraction of their expected ∼ 294 total (see Methods). Consistent with undersampling effects, neurons with more detected inputs showed stronger alignment between predicted and measured preferences (Supp. Fig. S1a). Despite these limitations, excitatory input alone was sufficient to predict postsynaptic preferred orientation.

**FIG. 2.**
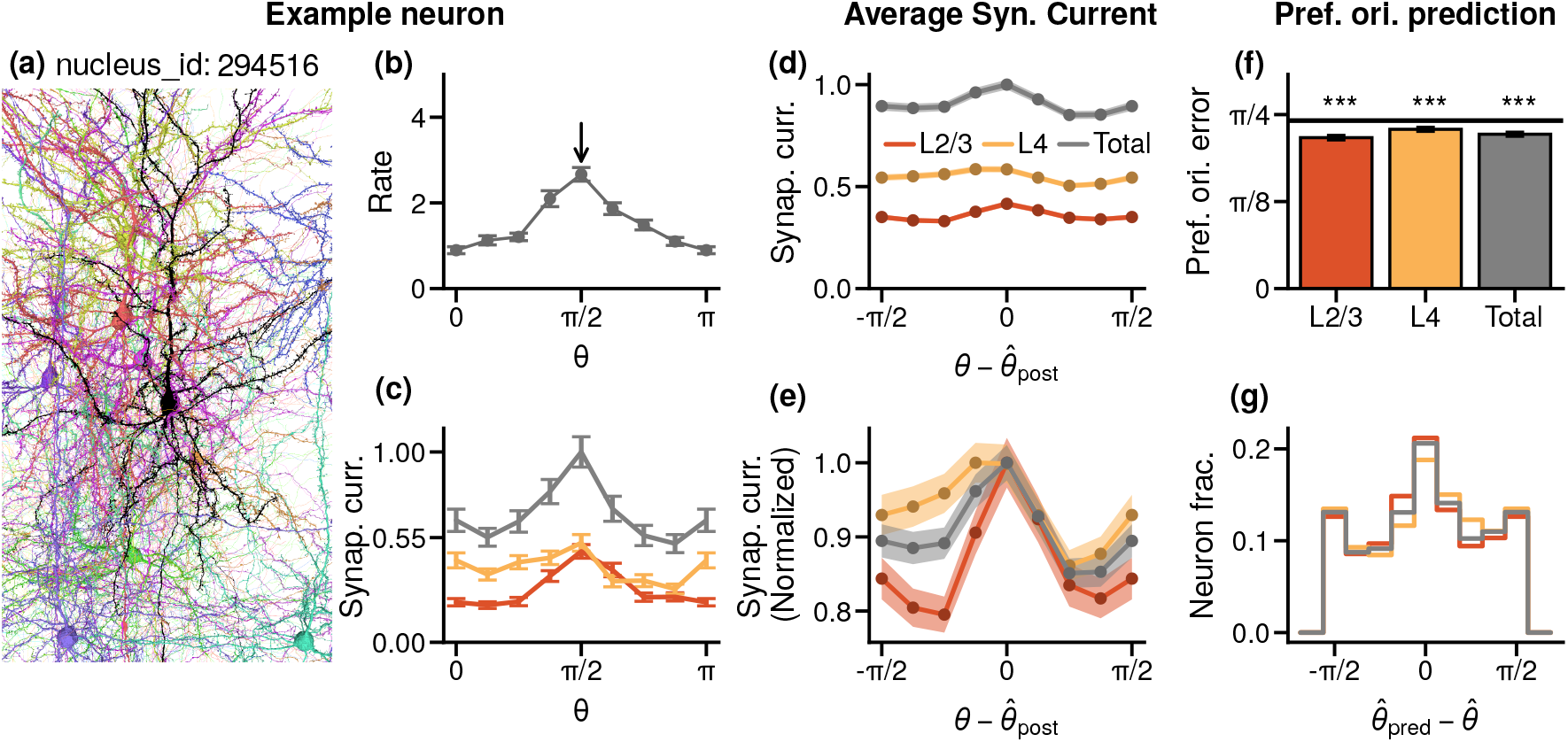
Feedforward and recurrent inputs provide comparable orientation-tuned drive to L2/3 neurons. **(a-c)** Synaptic inputs to an example L2/3 excitatory neuron (nucleus ID: 294516). **(a)** Neuroglancer EM reconstruction showing the postsynaptic neuron (black) and its presynaptic partners from L2/3 and L4. **(b)** Orientation tuning curve of the postsynaptic neuron (preferred orientation: *π/*2 rad, arrow). **(c)** excitatory synaptic currents from L2/3 (red) and L4 (orange) presynaptic populations, and their total (gray). Shading indicates the SEM. **(d**,**e)** Population-averaged synaptic inputs aligned to the postsynaptic preferred orientation (3609 L2/3 neurons; 256 presynaptic L2/3 neurons; 357 L4 presynaptic neurons). Here and in the following panels, shading indicates the SEM across neurons. Panel **(e)** shows the same as (d) but with currents individually normalized to their respective peaks. **(f**,**g)** Differences between each neuron’s measured preferred orientation (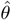) and that predicted from synaptic currents (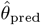). **(f)** Average difference. The horizontal line stands for a control result where preferred orientations of the postsynaptic neurons are randomly reshuffled. Asterisks denote significant differences from chance (bootstrap test, *p <* 0.05). **(g)** Distribution of 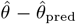for predictions based on total (gray), L2/3 (red), and L4 (orange) excitatory inputs.

We next examined the laminar structure of excitatory drive by separating inputs from L2/3 and L4 based on their presynaptic origin. The neuron in Fig. 2a-c, for example, received 30 excitatory inputs, with 15 from L2/3 and 15 from L4. Inputs from both layers were similar in magnitude and tuning. Extending this analysis to all orientation-selective L2/3 excitatory neurons revealed that, on average, 3.70 ± 0.06 inputs originated from L2/3 and 4.13 ± 0.06 from L4 (see Methods). Across the population, both sources showed comparable strength and orientation modulation, with L2/3 inputs marginally more tuned (Fig. 2d,e). Inputs from both layers predicted the postsynaptic preferred orientation (Fig. 2f,g), with L2/3 showing a significantly smaller average prediction error (0.68 ± 0.01 rad) than L4 (0.72 ± 0.01 rad, *p <* 5 *×* 10^−3^). Together, these results indicate that feedforward and recurrent excitatory currents are similarly tuned, providing comparable orientation-specific drive to L2/3 neurons.

### Synaptic current statistics suggest a key role for randomness in shaping selectivity

Neuronal selectivity depends on the composition of synaptic inputs, yet whether this arises from structured, tuningdependent connectivity or from random wiring remains unclear. Theoretical models provide a framework for distinguishing these possibilities. In structured models, each neuron is assigned a target preferred orientation (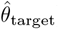) that guides connectivity, for example through “like-to-like” rules in which neurons with similar targets are more likely to be connected. The preferred orientation emerging from network dynamics (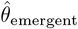) then matches the target, as connection randomness is assumed to be negligible [1, 18–2 In contrast, biological circuits inevitably contain substantial stochastic variability in connectivity, which can cause 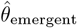 to deviate from 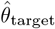. In balanced network models [16, 17], such variability is not merely noise but a key determinant of selectivity. Analyzing the statistics of synaptic inputs from the data therefore provides a principled approach to assess the relative contributions of structured and random connectivity to the emergence of selectivity. To test whether selectivity reflects structured or random connectivity, we analyzed the statistics of excitatory synaptic currents onto postsynaptic neurons. Most inputs originated from neurons with preferred orientations similar to the postsynaptic cell (Fig. 3a). This pattern could not be explained solely by an overrepresentation of specific orientations in the presynaptic population (Fig. 1b), since connection probability, independent of population bias, increased between neurons with similar orientation preferences (Fig. 3b). Nevertheless, a large fraction of inputs (83%) arose from neurons with dissimilar or no orientation tuning, and synaptic current strengths were highly heterogeneous, showing no systematic relationship with orientation difference (Fig. 3c).

**FIG. 3.**
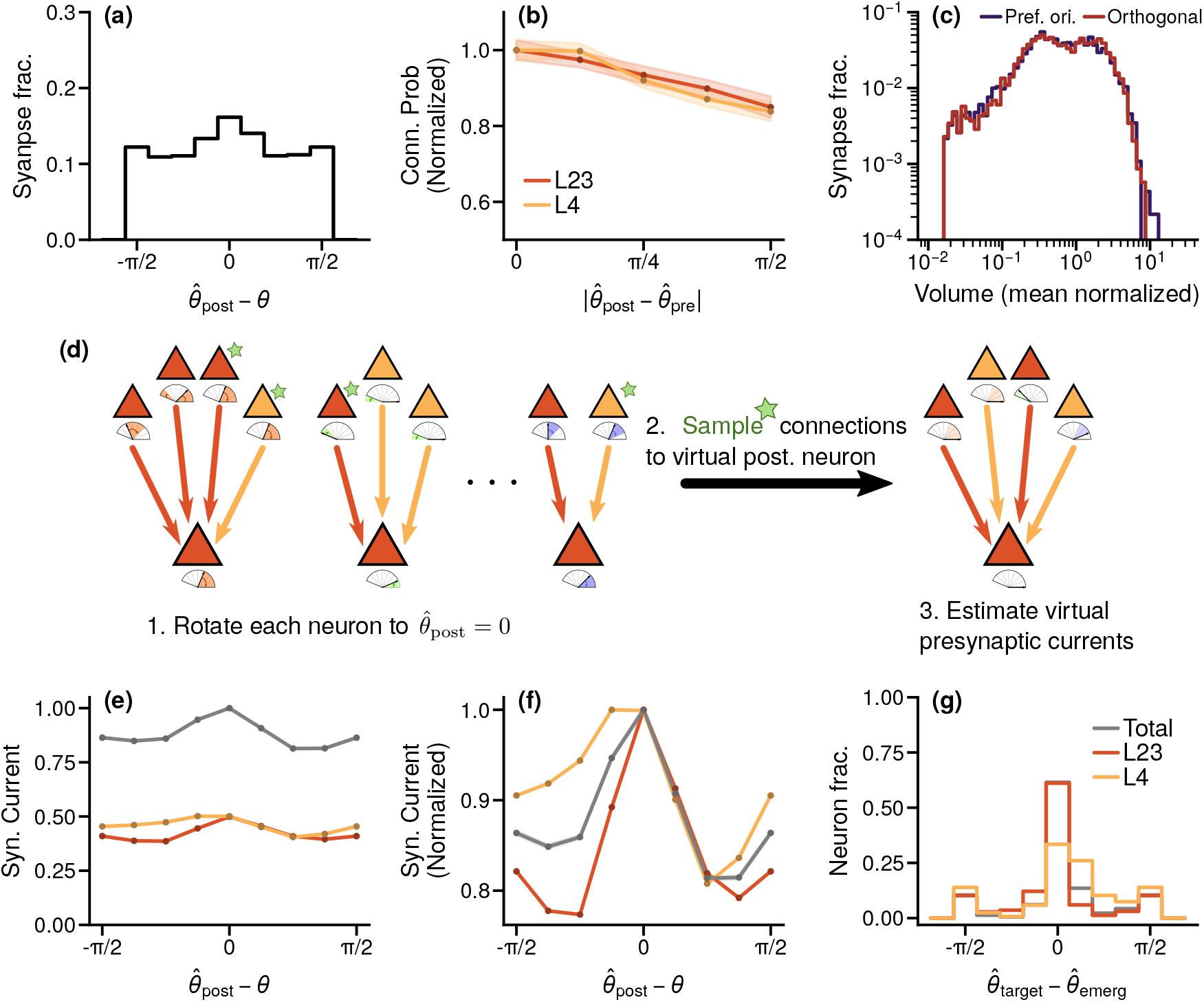
Synaptic input statistics reveal a dominant contribution of randomness to selectivity. **(a)** Fraction of inputs as a function of the difference between presynaptic and postsynaptic preferred orientations. **(b)** Connection probability as a function of orientation difference between presynaptic and postsynaptic neurons. Here and in subsequent panels, shading indicates the SEM. **(c)** Distribution of synaptic strengths as a function of orientation difference. **(d)** Schematic of the synthetic postsynaptic neuron model. Each model neuron was assigned a target preferred orientation (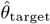) sampled from the empirical distribution (Fig. 1b). We selected a representative number of presynaptic partners while preserving the empirical ratio of L2/3 to L4 inputs and the observed connection probabilities as a function of presynaptic selectivity and preferred orientation. Presynaptic tuning curves were aligned to 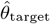, their currents summed, and the emergent preferred orientation 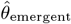 computed. **(e)** Average input currents for synthetic neurons. Currents from L2/3 (red) and L4 (orange) are normalized to the peak of the total input. **(f)** Same as (e) but normalized to each current’s individual peak. **(g)** Alignment between target and emergent preferred orientations for synthetic neurons, with the latter computed using all inputs (gray) or using only L2/3 (red) or L4 (orange) inputs. The target orientation was correctly predicted for 61% of the postsynaptic model neurons.

We next quantified how structure and variability in synaptic inputs influences selectivity by constructing a population of synthetic postsynaptic neurons (Fig. 3d). Following structured-model frameworks [1, 18–21], each neuron was assigned a target preferred orientation (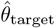) drawn from the empirical distribution (Fig. 1b). For each synthetic neuron, we randomly sampled a representative number of presynaptic partners from the pooled synaptic population (see Methods), preserving the observed L2/3-to-L4 ratio and the empirical dependence of connection probability on presynaptic selectivity and orientation (Fig. 3b). The emergent preferred orientation (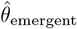) was then obtained by summing the resulting presynaptic currents. Excitatory inputs from L2/3 and L4 were comparable in magnitude and tuning (Fig. 3e,f), with L2/3 currents slightly weaker but more selective, consistent with empirical data (Fig. 2). Unlike the direct connectome-based reconstruction used in Fig. 2, this approach is not affected by undersampling, confirming that the laminar trends observed in the data are robust. The results were consistent across a physiologically relevant range of number of presynaptic partners (Supplementary Fig. S1b).

If structured connectivity dominates, the target and the emergent preferred orientations should closely align at the single-neuron level, whereas if randomness dominates, this alignment should be weak. In the synthetic population, the distribution of orientation differences (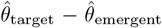) was centered near zero and significantly narrower than expected by chance (Fig. 3g), indicating that structure in connectivity contributes to selectivity. However, the distribution remained broad, with substantial variability. The average mean square error of the prediction was 0.38 (*∼ π/*8) rad, revealing that random fluctuations in synaptic inputs play a major role. Across synthetic neurons, L2/3 and L4 inputs showed comparable orientation tuning (Fig. 3e,f) and peak alignment with 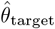 (Fig. 3g), with L2/3 inputs being slightly more tuned and predictive.

Together, these results indicate that selectivity emerges from the combined effects of structured connectivity and random synaptic variability, with feedforward and recurrent inputs contributing similarly. The interplay between structure and randomness may be further shaped by recurrent interactions, a possibility not captured by the synthetic neuron approach, which lacks network dynamics. To address this, we next model how excitation, inhibition, and network dynamics, constrained by the MICrONS dataset, reshape tuning and influence the alignment between 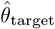and 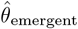.

### Connectome-constrained recurrent network models capture neural responses

To investigate how recurrent interactions shape selectivity in L2/3, we analyzed the dynamics of recurrent network models described by Eq. (1), constrained by structural and functional data from the MICrONS dataset [36].

A key challenge in constraining the network models was the incomplete specification of the connection matrices (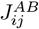). In addition to unknown global scaling factors that strongly influence network dynamics [56, 57, 62], the structural data contain missing synapses due to segmentation, reconstruction, and coregistration limitations. To overcome this, we modeled connectivity based on statistical properties inferred from the MICrONS dataset rather than individual connections. This approach follows theoretical work showing that cortical dynamics depend primarily on the statistical structure of connectivity rather than precise neuron-to-neuron wiring [56, 57, 62–6 We thus treated the reconstructed network as a single realization of a probabilistic ensemble in which connection probability and synaptic strength depend on pre- and postsynaptic cell type and orientation selectivity, with these relationships estimated directly from the data.

In the data, the probability of connection between L2/3 and L4 excitatory neurons decreased with increasing difference in preferred orientation (Fig. 3b). To incorporate this relationship, each neuron in the model was assigned a target preferred orientation, and connections were drawn probabilistically based on the orientation difference between pre- and postsynaptic targets (see Methods). This rule was applied to both recurrent and feedforward excitatory connections. Inhibitory connections were instead assumed to be random; modeling them with the same orientation-dependent rule as excitatory connections yielded qualitatively similar results (Supp. Figs. S3, S4, S5). Synaptic strengths were sampled from the empirical size distribution. Single-neuron responses were modeled using a validated approximation of the input–output function of leaky integrate-and-fire neurons driven by Gaussian noise [56–58, 67–7 with gain and offset parameters fit separately for excitatory and inhibitory cells and treated as free parameters.

Considering both single-neuron and network properties, the model contained eight free parameters. These were infered using simulation-based inference [71] to reproduce key response features of L2/3 excitatory neurons, including the summary statistics (value of the tuning curve at 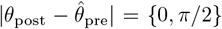; circular variance of the tuning curve; mean and variance of the logarithm of the rates; mean, variance and skewness of the circular variance; decrease of the connection probability at 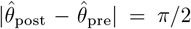) of the firing rates and circularvariance selectivity, the average tuning curve and the decrease in connection probability as a function of orientation difference (see Methods). This procedure yielded a family of solutions that captured these experimental features (Fig. 4a–e) with biologically plausible parameter values (Supplementary Fig. S2).

**FIG. 4.**
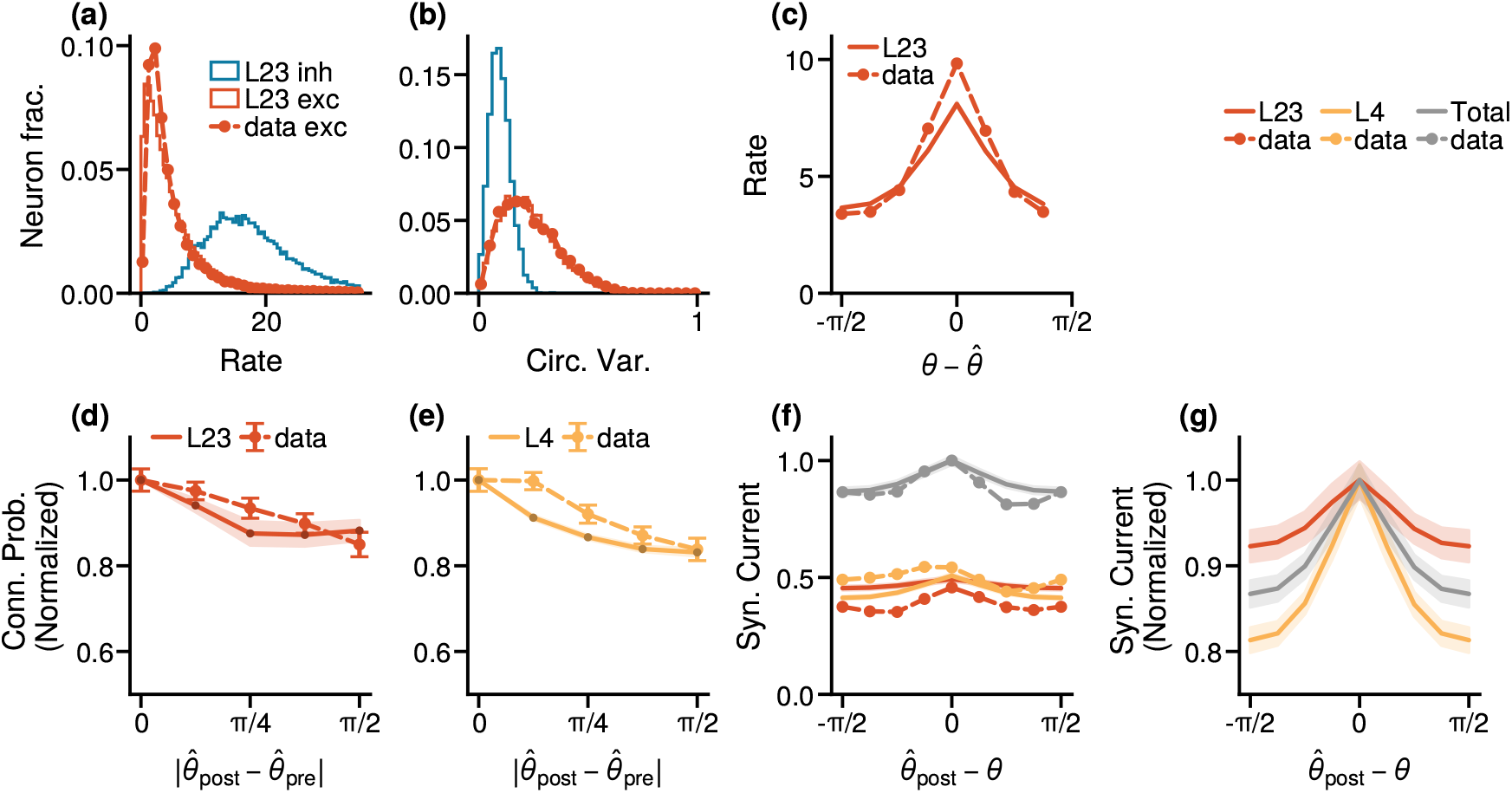
Connectome-constrained models capture diverse functional and structural features of L2/3. Comparison between observables measured in the network model (solid lines) and in the experimental data (dashed lines with markers). Error bars represent the SEM. **(a)** Firing-rate distributions for excitatory (red) and inhibitory (blue) neurons. **(b)** Circular-variance distributions for excitatory (red) and inhibitory (blue) neurons. **(c)** Average tuning curves of excitatory neurons in the model and data. **(d)** Connection probability between excitatory neurons within L2/3 as a function of the difference in their emergent preferred orientations. **(e)** Connection probability between excitatory neurons from L4 to L2/3 as a function of the difference in preferred orientations. **(f)** Excitatory synaptic currents (feedforward from L4, recurrent from L2/3, and total) as a function of the difference between the postsynaptic neuron’s preferred orientation (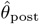) and the stimulus orientation (*θ*). Same as (f), but peak-normalized to highlight the relative tuning of feedforward and recurrent inputs.

The inferred models also reproduced several properties of the data that were not explicitly constrained. Despite optimizing only summary statistics, they captured the full distributions of mean firing rates and circular variances. Inhibitory neurons showed higher firing rates and lower selectivity than excitatory neurons (Fig. 4a,c), consistent with previous reports [49]. Blocking inhibition in the model caused runaway excitation, indicating that the network operates in an inhibition-stabilized regime where strong recurrent excitation is balanced by feedback inhibition, as observed in mouse cortex [72]. Finally, the tuning of excitatory inputs closely matched the data, with L2/3 and L4 inputs of similar strength and selectivity, although tuning in L4 was slightly stronger (Fig. 4f,g).

To further examine laminar contributions to tuning, we tested how well excitatory inputs from L2/3 and L4 predicted each neuron’s preferred orientation, replicating the analysis of Fig. 2g in the model. Consistent with the model’s excitatory tuning, L4 inputs were better predictors of postsynaptic preference (L2/3 error 0.59 rad; L4 0.40 rad, a 33% decrease, *p <* 10^−4^), opposite to the trend in the data (L2/3 error 0.68 rad; L4 0.72 rad, a 5% increase, *p <* 10^−4^). This difference diminished after subsampling model inputs to match the number of reconstructed synapses (L2/3 error 0.74 rad; L4 0.71 rad, a 4% decrease, *p <* 10^−4^). Remaining discrepancies likely reflect additional structure in the biological circuit not captured by the model, which may enhance selectivity in L2/3 inputs.

Together, these analyses show that the connectome-constrained model quantitatively reproduces the main features of the data, suggesting that similar mechanisms underlie orientation selectivity in both the model and the biological circuit. We next used the fitted model to perform *in silico* experiments to identify the circuit mechanisms driving selectivity in L2/3.

### Random connectivity dominates the emergence of selectivity

In the recurrent network models, structured connectivity followed a like-to-like rule based on each neuron’s target preferred orientation. During simulations, neurons developed emergent tuning and preferred orientations through their collective dynamics. Because connectivity included random components, the two orientations could differ. To quantify this, we measured the distribution of differences between target and emergent orientations in the best-fitting models. On average, the emergent orientation matched the target (Fig. 5a), but many neurons showed large deviations, indicating that random variability strongly influences selectivity. Compared with the non-recurrent, single-neuron framework (Fig. 3g), recurrent interactions did not reduce randomness and instead slightly amplified it, increasing the mean square error to 0.51 rad (a 35% rise).

**FIG. 5.**
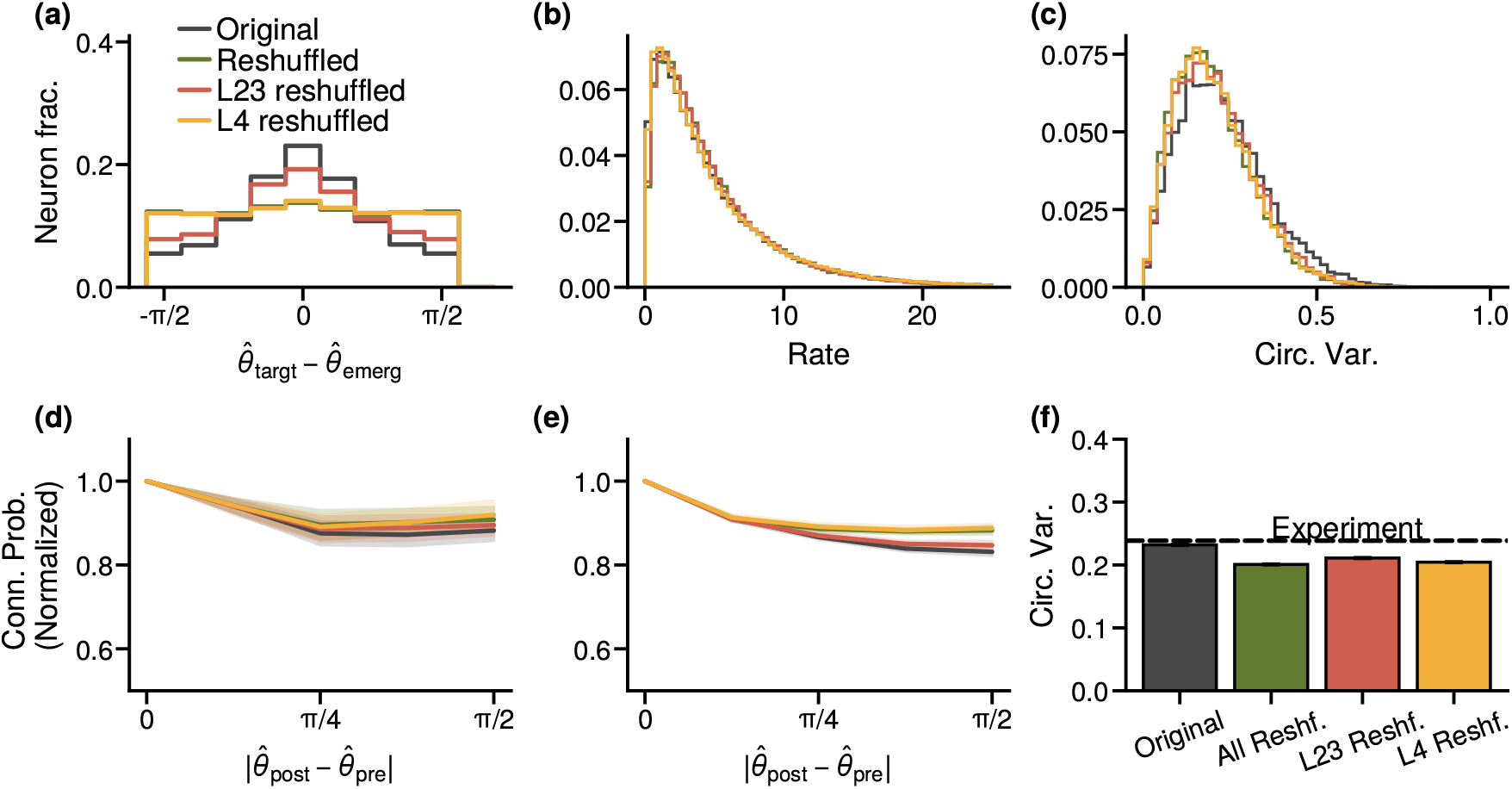
Connectivity randomization reveals the respective roles of structured and random connections in shaping selectivity. **(a)** Distribution of differences between target and emergent preferred orientations (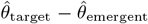) in recurrent network models. Here and in the following panels, results are shown for the best-fit structured model (black) and for models with randomized feedforward (orange), recurrent (red), or all (green) connections. **(b)** Firing-rate distributions of excitatory neurons across model variants (best-fit vs. shuffled). **(c)** Circular-variance distributions under different randomization conditions. **(d)** Connection probability between L2/3 excitatory neurons as a function of emergent preferred-orientation difference. **(e)** Connection probability from L4 to L2/3 neurons as a function of emergent preferred-orientation difference. **(f)** Mean circular variance across the 100 best-fit models under normal conditions (black) and for each randomization scenario (colors as in panels above). Error bars represent the SEM.

To assess the respective roles of structured and random connectivity in shaping selectivity, we shuffled the model’s connections while keeping all parameters fixed. This preserved global statistics such as connection density and synaptic strength but removed orientation-dependent structure. As expected, shuffling disrupted the alignment between target and emergent orientations (Fig. 5a). Remarkably, however, it produced only minor changes in network responses (Fig. 5a–f), leaving the mean firing rate unchanged (Fig. 5b) and causing a weak but significant reduction in selectivity (Fig. 5c,f). Disrupting connectivity structure also had only a weak effect on the like-to-like connection probability as a function of emergent preferred orientation (Fig. 5d). This finding is consistent with previous theoretical work showing that functional specificity can emerge even in randomly connected networks due to statistical heterogeneity in the generative connectivity rule [17].

Although shuffling had only a modest impact on network responses, it produced a measurable reduction in selectivity, indicating that structured connectivity acts as an amplifier that enhances a baseline of selectivity largely generated by random connections. Previous theoretical work [19–21] showed that like-to-like connectivity sharpens tuning, consistent with our finding that it decreased slightly after shuffling (Fig. 5d,e). Feedforward and recurrent structures contributed differently to this process: shuffling feedforward connections alone impaired the alignment between target and emergent preferred orientations and reduced like-to-like connectivity to a similar extent as shuffling all connections (Fig. 5a,d,e), whereas shuffling recurrent connectivity had a smaller effect. Likewise, feedforward shuffling reduced orientation selectivity, comparable to the reduction observed when all connections were shuffled, while recurrent shuffling caused a weaker decrease (Fig. 5c,f).

Together, these results support an amplification mechanism in which structured feedforward inputs from L4 align target and emergent orientations, enabling recurrent like-to-like connectivity within L2/3 to further enhance selectivity.

In summary, these analyses show that randomness in connectivity is the primary source of selectivity in L2/3, establishing a baseline of orientation tuning, while structured connectivity serves as a secondary mechanism that refines and amplifies it.

## DISCUSSION

Feature selectivity is a defining property of cortical computation [1–15], yet its circuit origins remain debated [1, 16–21]. To address this, we combined largescale functional recordings with a synapse-resolution connectome of mouse visual cortex from the MICrONS dataset [36] to build mechanistic models linking circuit structure to orientation selectivity in layer 2/3 of V1. We first analyzed single neurons in the MICrONS dataset [36] and found that feedforward input from layer 4 and recurrent input within layer 2/3 contribute comparably to selectivity. Analyses of synthetic neurons constructed from empirically measured input statistics confirmed this and showed that variability in connectivity is a key determinant of selectivity. We then combined functional and structural data with parameter inference to build recurrent network models that quantitatively reproduced both the response properties and connectivity patterns observed in the data. These models revealed that randomness in connectivity, rather than precise wiring, plays the dominant role in generating selectivity, while structured inputs provide a secondary, amplifying effect.

Several theoretical models have been proposed to explain orientation selectivity, invoking either structured feedforward [1, 18] or recurrent [19–21] connectivity, or unstructured networks with strong recurrent interactions operating in an inhibition-dominated regime [16, 17]. Our results indicate that the latter provides the dominant mechanism underlying orientation selectivity in layer 2/3 of mouse V1. Nevertheless, the emerging picture is more nuanced. We observed “like-to-like” feedforward and recurrent connectivity, consistent with previous studies [24–28] and with assumptions of structured models [1, 18–21]. Although most selectivity arises independently of this structure, it plays an important amplifying role. Structured feedforward inputs bias emergent preferred orientations toward those of presynaptic partners, while recurrent connectivity enhances excitation among similarly tuned neurons, jointly strengthening population-level selectivity.

Our analysis of the MICrONS dataset [36] showed that connection probability in both L2/3 and L4 decreases with increasing difference in preferred orientation and that tuned neurons frequently receive inputs from non-selective partners, consistent with prior physiological and anatomical studies [24–28, 73]. An independent analysis of the MICrONS dataset by Ding et al. [52] reported no significant like-to-like dependence among excitatory connections within L2/3, likely due to methodological differences. Ding et al. estimated preferred orientation using digital-twin predictions, whereas we derived it directly from visually evoked responses to parametric stimuli. Although digital-twin models perform well on average, they can deviate at the single-neuron level [74], adding noise to orientation estimates. In addition, their analysis used a smaller and more heterogeneous sample (148 presynaptic and 4,811 postsynaptic neurons across layers and areas), while ours focused on a larger, layer-specific population (611 presynaptic and 4,254 postsynaptic neurons exclusively in L2/3 and L4 of V1). This difference may arise from their stricter proofreading criteria, whereas we included neurons with unproofread dendrites, which are typically well segmented, and at least clean proofread axons. Because reconstruction errors are expected to be independent of neuronal tuning, this inclusion should not bias orientation-dependent connectivity. Consistent with this expectation, we repeated the analysis across subsets with different proofreading status and found that proofreading primarily affected the absolute connection probability, while the decrease in connectivity with increasing difference in preferred orientation remained robust (Supp. Fig. S6). The agreement of our results with previous physiological and anatomical findings [24–28, 73] further supports the validity of our approach.

Connectomic datasets provide unprecedented insight into cortical circuitry, but current reconstructions, including MICrONS [36], remain incomplete. Automatic segmentation introduces reconstruction errors [37, 75], and while manual proofreading improves accuracy, it is impractical at cortical scale and automated corrections are still developing [48]. Moreover, functional properties such as synaptic strength, neuronal excitability, and intrinsic dynamics cannot be inferred from structure alone. As a result, even state-of-the-art connectomes offer only a partial view of circuit wiring. These limitations raise a key question: how can incomplete connectomic data be used to uncover circuit mechanisms? Our approach addresses this challenge by integrating structural and functional information within a theoretical framework that focuses on population-level statistics rather than individual connections. This strategy builds on theoretical work showing that in highly connected cortical circuits, collective dynamics depend mainly on statistical connectivity patterns rather than precise synaptic details [62–66]. Leveraging this regime, we extracted connectivity statistics from the partial reconstruction, generated synthetic networks consistent with these statistics, and used Bayesian inference on functional data to estimate unobserved quantities such as input–output relationships and average synaptic strengths. This data-constrained modeling reproduced key experimental observations and, through in silico perturbations, revealed the circuit mechanisms underlying orientation selectivity. More broadly, it shows how combining structural and functional constraints within a principled theoretical framework can turn incomplete connectomes into powerful tools for mechanistic discovery.

The advent of large-scale connectomic reconstructions marks a major advance in neuroscience, providing unprecedented access to the circuit basis of neural computation. Complete wiring diagrams have long been available for species with smaller brains, such as zebrafish and *Drosophila*, and have been successfully integrated into theoretical and mechanistic models [38–47]. Cortical connectomes, however, are only now reaching the scale needed for detailed modeling of mammalian circuits [26, 36, 76], and general strategies for integrating these data into mechanistic models remain scarce. Our framework bridges this gap by combining structural and functional information within a principled theoretical context to infer circuit mechanisms from incomplete data. Applied to layer 2/3 of mouse visual cortex, it revealed how random connectivity can generate orientation selectivity, while structured inputs refine and amplify it. More broadly, this approach offers a general path for linking structure and function across brain regions and species and will become increasingly valuable as connectomic reconstructions expand to larger and more complex neural systems.

## MATERIALS AND METHODS

### Analysis of the MICrONS dataset

We relied on the MICrONS dataset [36] v.1300 to extract functional and structural properties of neurons in layers 2/3 and 4 of mouse V1.

We first filtered the units of interest in this region. We extracted information about area assignment from `nucleus_functional_area_assignment` table provided in v1300 of the data and filtered for V1 units.

Layer assignment was based on AIBS neuron classification from the table `allen_column_mtypes_v2` [77]. The classification contains information about the different morphologies of pyramidal neurons in each layer and inhibitory subtypes. We discretized pial depth in small bands of 10 µm and assigned a layer to each band from the most common type of excitatory neuron in the band. For example, a band with majority of 23P excitatory neurons and presence of interneurons will be assigned L2/3. Our layer boundaries are compatible with previous results [52] and are adequate for the recent v1300 version of the data. L2/3 spans from 70 µm to 250 µm, and L4 from 250 µm to 360 µm).

#### Functional properties

We extracted responses from neurons in layers 2/3 and 4 of V1 from the functional dataset. Neurons in these areas were identified using table `coregistration_manual` v4. We identified 7759 functionally matched excitatory neurons, of which 4620 were in L2/3 and 3139 were in L4. Of these, 256 had proofread (`clean` or `extended`, as defined in the next section) axons in L2/3 and 357 in L4.

We analyzed trials with ‘Monet’ stimuli—parametric textures that vary in orientation and spatial frequency—each trial presenting one of 8 orientations from 0 to 180 degrees, with drift perpendicular to the stimulus’s orientation (details in [36]). To characterize the selectivity of each neuron to stimulus orientation, we first identified the frames in each recording session during which the animal was exposed to a specific stimulus orientation. We then calculated the average response and its standard error across these repeated trials for all orientations using the deconvolved spike traces. These responses were fitted with a von Mises function [78] utilizing least square optimization through the `curve_fit` function implemented as part of `scipy` library in Python,

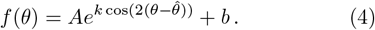

Here, *A* is the amplitude of the cosine wave, *k* is a width parameter, 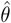 is the preferred orientation and *b* is a baseline rate. Although the fit returns a continuous estimation for 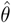, we round the values to the 8 possible experimental orientations in order to compare directly with the data.

We consider a fit successful when its *R*^2^ coefficient is larger than 0.5. Neurons with a smaller value are considered to be non-selective. For succesful fits, we compute the trial measurements of average firing rate at the preferred orientation 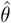 and the orthogonal orientation 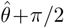. We then performed a Wilcoxon signed-rank test on the two sets; a p-value *<*0.01 indicated significantly higher activity at 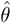, classifying the neuron as selective. If the test was not significant, we also classified the neuron as non-selective. All p-values were corrected to set a false discovery rate of 5%.

We characterized the tuning of selective neurons through circular variance

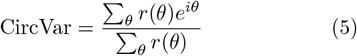

where *i* is the imaginary constant, and the sum goes over the potential discrete values of the preferred orientation.

This parameter is close to 0 for non selective neurons and close to 1 for highly selective ones.

#### Structural properties

We characterized connectivity within layers 2/3 and 4 of V1 using version v1300 of the structural dataset. Neuronal layers were identified based on depth as described above.

Through this approach, we identified a total of 24774 neurons, of which 22647 are excitatory (9487 L4 and 13160 in L2/3) and 2127 inhibitory neurons, of which 1299 are in L2/3. Neurons with axonal proofreading had smaller numbers: a total of 972 neurons, of which 453 were in L2/3. Synapses between the neurons were obtained through API calls to the synapse table to query connections of the neuron subset of interest.

As discussed in the main text and in [48], automatic synaptic connection reconstructions are subject to error, which can be addressed through manual proofreading. In the MICrONS dataset, dendritic and axonal reconstructions were proofread independently, with different levels of precision. Proofreading efforts have followed different strategies and labelling which have changed during the development of the MICrONS project. In the case of axons, partial proofreading indicates that a branch was followed until its termination. For fully extended proofreading, all end points of the axons are manually checked and re-extended if possible [61]. In this paper, we refer to partial proofreading as `clean` axons, and extended as `extended`, in accordance with a previous nomenclature adopted by the MICrONS team. It is important to remark that in our case, in v1300, no other strategies (such as truncated or interarea) were present. In our analysis, neurons were selected based on their reconstruction status, filtering for manual proofreading in axonal reconstructions due to the higher error rates associated with their smaller size. Dendrites, being thicker, are generally well-segmented, with most merge errors involving small axons [79]. We extracted connectivity properties from synaptically connected pairs with at least `clean` axonal proofreading and no dendritic proofreading. We estimated connection strength using synapse size (cleft volume in voxels) from the table of synapses; when multiple synapses were present, we combined their sizes to measure the total connection strength. Synaptic sizes were then normalized such that the average size was set to 1.

The data analysis pipeline is fully automated, documented and easy to use, and will be released as open source code upon the publication of the final version of the paper. Part of the pipeline is already public and available on GitHub(https://github.com/MICrONS-Milano-CoLab/MICrONS-datacleaner/) as an independent package. The codebase also contains several utilities to work with the processed data, to easily filter neurons and synapses for common criteria, work with neuronal activity, etc.

### Inference of network statistics

#### Connection probability and strength

To determine the statistics of connectivity between different neuron types, we cataloged all excitatory and inhibitory neurons in L2/3, as well as excitatory neurons in L4 (see above for determination of excitatory and inhibitory neurons). From this list, we extracted all recorded synapses between these neurons to construct a network. We do not consider individual synapses. When two neurons are linked through multiple synapses, we consider a single link with an effective size equal to the sum of all synaptic volumes. Thus, the resulting network is directed and weighted and can be represented with a connectivity matrix. We quantified connection probability between each pair of presynaptic and postsynaptic neuron types and selectivity properties by calculating the fraction of nonzero entries in this matrix relative to the total number of possible connections between the selected populations.

#### Estimation of the in-degree

We estimated the indegree of excitatory neurons by combining the connection probability between excitatory pairs with the total number of excitatory neurons. Connection probability was measured from neuron pairs whose axons were proofread to the `extended` level and dendrites to the `clean` level to ensure accuracy. The total number of excitatory neurons was obtained by counting all identified excitatory (E) cells in the dataset. This approach yielded an expected in-degree of *K*_*EE*_ =168. Repeating the calculation for axons with at least clean proofreading gave 95. For most analyses, we therefore set *K*_*EE*_ = 150, consistent with these estimates. From the probabilities of connection it is possible to estimate the total excitatory input from both layers 2/3 and 4 grows up to be *K* = 293. To provide bounds, we also computed a lower and an upper limit for the in-degree. The lower bound (*K*_*EE*_ = 23, *K* = 45) was obtained by considering only inputs to L2/3 excitatory neurons from other objects explicitly identified as neurons, using `extended` proofreading for both axons and dendrites. The upper bound (*K* = 1305) was derived from synapse-level data. Because two neurons can form multiple synapses, we first estimated the average number of synapses per connected pair using `extended` proofreading, then used the total synapse count per postsynaptic neuron, which is reliable in the dataset, to estimate the total number of inputs each neuron receives across all areas.

### Analyses of selectivity of synaptic currents

#### Direct derivation of synaptic currents from connectome

To probe the relationship between postsynaptic firing rates and synaptic currents as a function of stimulus orientation predicted by Eq. (3), we identified, for each selective postsynaptic neuron, all functionally matched presynaptic partners with at least a `clean` level of axonal proofreading. We calculated the individual synaptic current contributions by multiplying the tuning curve of each presynaptic neuron (i.e., its firing rate as a function of stimulus orientation) by the weight of the corresponding connection. The weight of each connection corresponds to the sum of all synapse sizes from the presynaptic neuron to the postsynaptic one. For non-selective presynaptic neurons, we modeled responses as orientation-independent by assigning a constant firing rate equal to the average across all stimulus orientations. Summing across all synapses provided an estimate of the total synaptic current for the postsynaptic neuron. The orientation at which this current was maximal predicted the neuron’s preferred orientation, which can be compared with that measured from its postsynaptic firing-rate tuning curve. This procedure yielded the results shown in Fig. 2. To assess the model’s efficacy at the population level, we repeated this procedure for all neurons in the dataset, measuring the difference between each neuron’s observed preferred orientation and the orientation predicted by synaptic currents. Fig. 2 presents a histogram of these differences. As a control, we shuffled the dataset by randomly assigning each postsynaptic neuron a set of presynaptic partners with randomly sized synapses, then computed synaptic currents as before.

#### Statistical derivation of synaptic currents from connectome

To investigate the statistics of synaptic currents in selective neurons, we construct model postsynaptic neurons with input statistics that resemble the ones of the original data. To do so, we compiled a list of all synapses to selective neurons along with the tuning curves of their corresponding pre- and postsynaptic neurons. Connections are filtered so presynaptic neurons had at least a `clean` level of proofreading at least. Then, we rotated each tuning curve so that the postsynaptic neuron was aligned to a zero orientation preference. Now it is possible to generate model neurons by randomly sampling synapses according in the table. We summed these inputs to calculate the synaptic current, following the same method as in the direct derivation of synaptic currents. For each model neuron, we extracted the contributions from both L2/3 and L4 components, repeating the procedure 1000 times. Fig. 3e,f displays the mean synaptic current across these repetitions, with error bars representing the standard error of the mean. Fig. 3g shows a histogram of the peak positions of these currents across repetitions.

### Simulation and analysis of recurrent network models

#### Mathematical description of the network model

Dynamics of neural activity in the network model is Eq. (1). The single-neuron input-output function was taken to be the f-I curve of leaky integrate-and-fire neurons driven by white noise [67]. For a neuron belonging to population *A* ∈ [*E, I*] in response to an input *µ*, this was given by

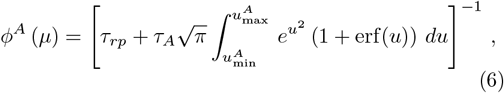

with

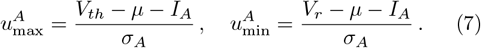

The parameters *σ*_*A*_ and *V*_*th*_ control the slope of the input-output function and were assumed to be independent of neuron type. The parameter *I*_*A*_ controls the offset of the input-output function and implicitly models feedforward inhibition from L4; this parameter was assumed to differ between *E* and *I* neurons. We fit *σ*_*A*_ and *I*_*A*_ to the data, while standard values were assumed for the refractory period *τ*_*rp*_ = 2 ms, resting potential *V*_*r*_ = 10 mV, and threshold *V*_*th*_ = 20 mV.

We modeled networks comprising a total of *N* = 3000 neurons, with 1583 excitatory (E) neurons, 241 inhibitory (I) neurons, and 1177 driving (X) neurons. The orientation-tuning responses of X neurons were sampled from the functional data, preserving the observed statistics for the fraction of selective neurons and the distribution of preferred orientations among selective neurons. We assumed that the reconstructed connectivity in MICrONS represents a single sample from an underlying probabilistic ensemble in which both connection probability and synaptic strength depend solely on the cell types and orientation selectivity of the pre- and postsynaptic neurons. These statistical dependencies were inferred directly from the data. In the structural dataset, the connection probability between L2/3 and L4 excitatory neurons decreased with increasing difference in preferred orientation (Fig. 3b), whereas synaptic strengths were broadly distributed and independent of orientation (Fig. 3c). To incorporate these empirically derived constraints into the model, each neuron was assigned a *target* preferred orientation drawn from the distribution observed in the functional dataset. Connections were then established probabilistically according to the orientation difference between pre- and postsynaptic neurons, paralleling structured network models of selectivity [1, 18–21]. Synaptic strengths were sampled from the empirical size distribution.

Because the observed orientation dependence of connectivity is defined in terms of *emergent* rather than target tuning, we parameterized connection probability as a function of target orientation difference and calibrated the parameters to reproduce the experimentally measured dependence on emergent differences. We applied this selectivity-dependent rule to both recurrent excitatory connections and feedforward connections from L4 (X) to L2/3 (E) neurons, such that the probability of connection between neurons of types *A* and *B* was given by

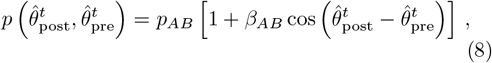

where 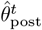 and 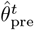 are the target preferred orientations of the post- and pre-synaptic neurons, respectively. Here, *p*_*AB*_ represents the baseline connection probability between neurons of types *A* and *B*, as obtained from the structural dataset, and *β*_*AB*_ modulates this probability according to selectivity of the neurons. In the data, we observe *β*_*EE*_ = 0.17 and *β*_*EX*_ = 0.15 for the emergent orientations. For the best fitted model, we have *β*_*EE*_ = 0.30 and *β*_*EX*_ = 0.15. The full estimated posterior distribution can be seen in Supp. Fig. S2 Thus, a slighly larger structural ‘like-to-like’ structure in the recurrent layer is required to reproduce the emergent connection probability of the data. In the case of simulations with tuned inhibition, we find *β*_*EE*_ = 0.14 and *β*_*EX*_ = 0.15. Since by construction the model is way more structured, the initial structural probability can be smaller, with values compatible with the observations in data.

#### Simulation procedure

We conducted simulations of the network dynamics in Python. For each parameter set and stimulus orientation *θ*, we generated ten realizations of the network structure, including the connectivity matrices 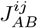 and the firing rates of the external population 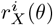. We simulated the network dynamics according to Eq. (1) using an explicit Euler method with timestep Δ*t* = 5 · 10^−3^ ms. We ran the simulations for a total duration of 200*τ*_*E*_ = 4 s and stored the computed firing rates every *τ*_*I*_*/*3 = 3.33 ms. We determined the firing rate of each cell by calculating its average rate over the simulation period, excluding the initial 10*τ*_*E*_ = 0.2 s. We repeated this procedure for each fixed value of *θ* and obtained tuning curves for the neurons by combining results across different values of *θ*.

#### Fitting procedure

The free parameters in the model are a global synaptic coupling *J*, the inhibitory coupling *g*, the inhibitory currents from L4 (*I*_*E*_ and *I*_*I*_), slope of the nonlinear input-output function *σ*_*E*_, *σ*_*I*_ and the modulation of the connection probability due to different preferred orientation *β*_*EE*_ and *β*_*EX*_. We identified the set of parameters that best fit the data using a simulation-based inference (SBI) approach employing the sbi package [71]. We conducted a total of 10^5^ simulations by randomly sampling 10^4^ parameter sets from a uniform prior and running 10 repetitions for each set, using different realizations of the connectivity matrix and L4 responses. The simulations were used to train a deep neural network via the SNPE algorithm [71] using the average tuning curves of excitatory neurons in L2/3 as summary statistics. The trained network can estimate a posterior distribution for the observed average tuning curves. We sampled another 10^5^ sets of parameters from the estimated posterior to run additional simulations. This approach allowed us to generate a large number of simulations with summary statistics close to the ones observed in the data. Finally, we looked for simulations from the posterior that jointly minimized the error of the average firing rate distribution, circular variance distribution, and connection probabilities in layers 2/3 and 4. To do so, we computed the mean square error in rate, connection probability and circular variance for each simulation and obtained the distribution of each error separately. Then, we found simulations that jointly minimized the three of them by getting those that were in the 5% percentile of each distribution. The results shown in Fig. 3 and Fig. 4 presents data from the simulations the provided the best overall simulation.

##### Analysis of model responses

For network models with parameter sets that produced responses matching experimental data, we performed further analysis to measure selectivity and determine the preferred orientation of each neuron. We then used the same network model and scrambled the columns of the E-E subnetwork, the E-X subnetwork, or both, running simulations on each scrambled configuration. In these simulations, L4 responses were identical, and only the connectivities were changed. These scrambles removed the tuning-dependent structure in connectivity. In each scrambled network, we remeasured neuronal selectivity to estimate the resulting circular variance. We also determined the preferred orientations and measured the connection probability as a function of differences in emergent preferred orientation, following the same procedure applied to experimental data.

## DATA AND CODE AVAILABILITY

Data analysis and network simulation codes are available at https://github.com/NeuroBLab/con-con-models. A general purpose package to work with the MICrONS dataset has been released at https://github.com/MICrONS-Milano-CoLab/MICrONS-datacleaner/. The data used in this study is publicly available [36].

## ACKNOWLEDGMENTS

V.B acknowledges funding by the NextGenerationEU, in the framework of the FAIR—Future Artificial Intelligence Research project (FAIR PE00000013—CUP B43C22000800006). Simulations were done at the PROTEUS cluster at the Carlos I Institute for Theoretical Physics, Granada, Spain, funded by the Spanish Ministry and Agencia Estatal de investigació;n (AEI) through Project I+D+i Ref. No. PID2020–113681 GB-I00, financed by MICIN/AEI/10.13039/501100011033

S0

## Supplementary information

## I. MECHANISTIC MODELS OF FEATURE SELECTIVITY IN CORTICAL CIRCUITS: CHALLENGES AND APPROACHES

This section provides the theoretical background for the discussion in the main text regarding the circuit mechanisms that can give rise to feature selectivity in recurrent cortical networks. Here, we formalize the problem introduced in the *Introduction* and summarize analytical results from three classical classes of models (feedforward, recurrent, and balanced networks) that have been proposed to explain how selectivity can emerge from underlying connectivity. We start by considering a general rate-based network model and derive under what conditions neurons develop selective responses to sensory stimuli.

We start from Eq. (1) of the main text, reproduced here for convenience:

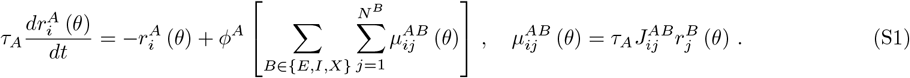

In this equation, *θ* represents the stimulus, which could correspond to the orientation of a grating, the position of a dark spot on a screen, or the light intensity at each pixel of an image. We assume that neurons in the input population *X* are selective to specific features of the stimulus. For simplicity, we assume that neurons differ only in their preferred feature:

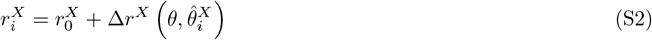

where 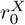 is the baseline activity, 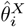 is the preferred feature of neuron *i*, and Δ*r*^*X*^ describes the modulation of the neural response to stimulus *θ*. For orientation selectivity, 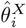 is the neuron’s preferred orientation and 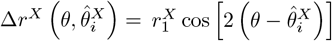. In the Hubel and Wiesel model of thalamic selectivity, *θ* = *θ* (*x, y*) is the luminance at each pixel, 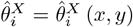 defines the receptive field (e.g., a difference-of-Gaussians ON-center/OFF-surround profile), and 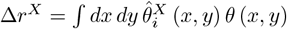.

Given the model above, what are the conditions under which *E* neurons develop selectivity, i.e., exhibit a modulation 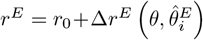 with Δ*r*^*E*^ = *O* (1)? We first show that this is not trivial and then analyze three classes of models that have been proposed to account for it. Throughout, we assume 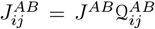, where Q is an Erdös–Rényi adjacency matrix with connection probability *q*.

A necessary condition for *E* neurons to develop feature selectivity is that their firing rates 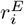 (and consequently their input currents 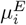) exhibit *O* (1) modulation with respect to *θ*. However, this is not trivial, because the feedforward input 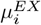 received by a neuron is expected to depend only weakly on *θ*. Indeed,

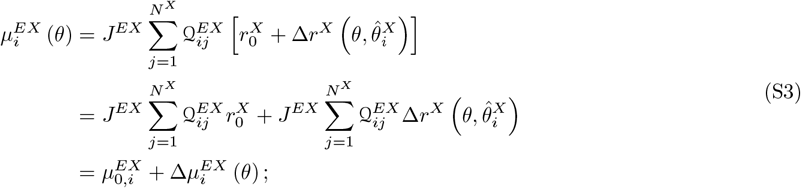

where 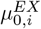 is the nonselective baseline term and 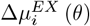 captures the stimulus-dependent modulation. Both can be approximated as Gaussian random variables with

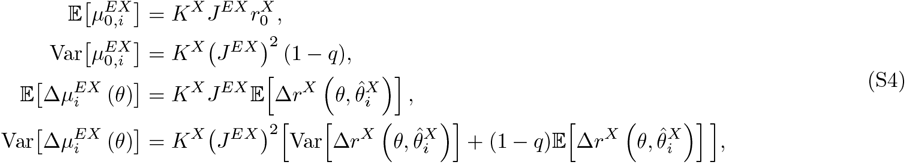

with *K*^*X*^ = *N*^*X*^*q* denoting the average in-degree from population *X*.

Since rates are *O* (1), the baseline component 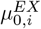 has a mean *O* (*KJ*) and neuron-to-neuron fluctuations 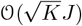 it does not depend on the stimulus and therefore cannot generate tuning. In contrast, the selective component 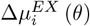 depends critically on 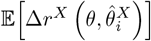, which vanishes if preferred features are uniformly distributed. For instance, in the case of orientation selectivity:

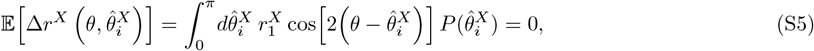

if 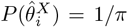. Thus, the selective input component scales as 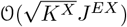, much smaller than the nonselective component *O* (*K*^*X*^*J*^*EX*^), with their ratio 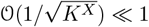.

How can *E* neurons develop strong selectivity in response to such weakly selective inputs? Three classical mechanisms have been proposed:

- **Feedforward models [1, 18]**. Structured feedforward connectivity breaks the assumptions of the central limit theorem by introducing correlations in the input patterns. Specifically, only *X* neurons with receptive fields aligned along a common axis have high probability of connecting to a given *E* neuron. The model assumes weak synapses *J* = *O* (1*/K*) to ensure that the mean input remains *O* (1). This architecture allows selectivity to be inherited from structured input correlations.
- **Recurrent models [19–21]**. Recurrent structure amplifies weakly selective feedforward inputs. Here, *E* neurons are assigned preferred features 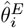, and neurons with similar tuning are more strongly connected, for example through 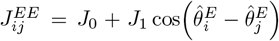. In this model, the selective component of the response 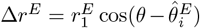 is amplified by a factor ∼ (2 − *J*_1_)^−1^ as *J*_1_ → 2, and for larger *J*_1_ the network can generate self-sustained bump states.
- **Balanced models [16, 17]**. These models assume random connectivity and strong synapses 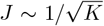. In this regime, large nonselective inputs 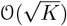 are dynamically canceled by inhibition, allowing the weaker *O* (1) stimulus-dependent component to determine the firing rates. Thus, selectivity arises not from structured wiring but from the balance between excitation and inhibition in random networks.

## II. SUPPLEMENTARY FIGURES

**FIG. S1.**
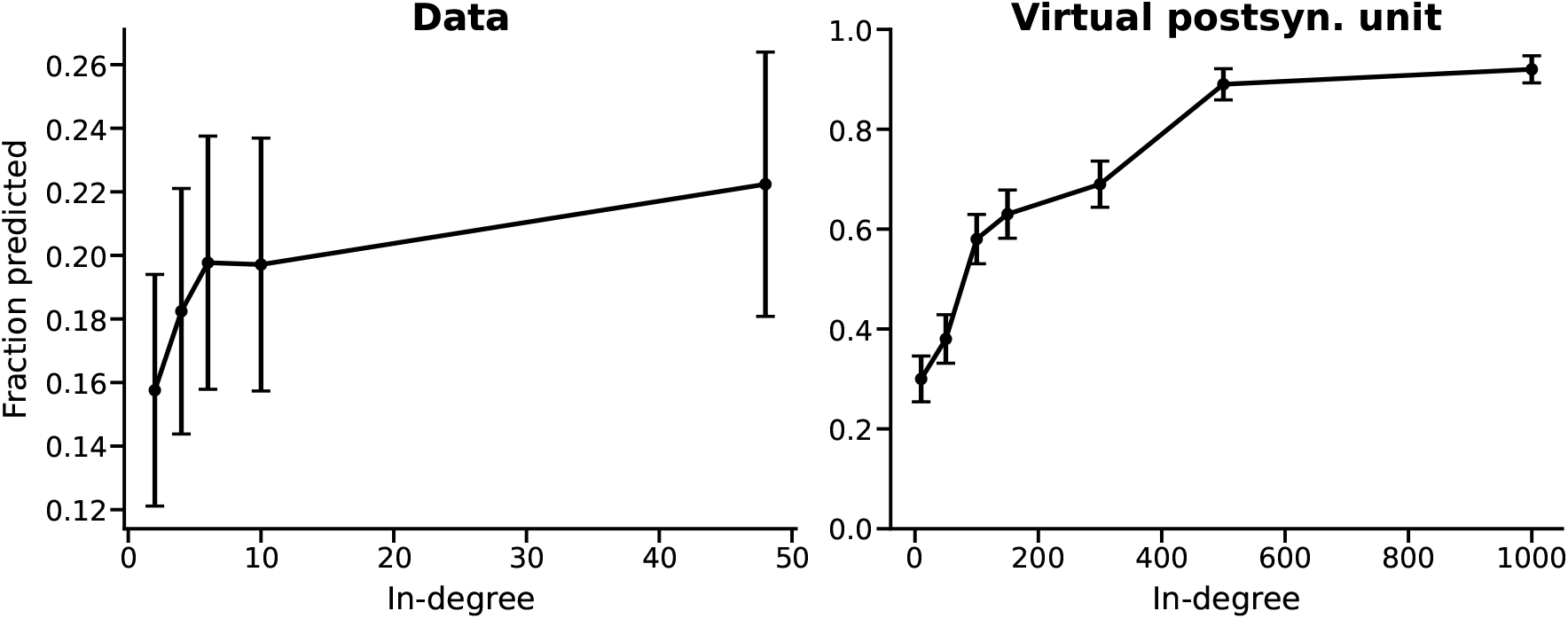
Supplementary analysis of the dependence on in-degree *K*. **(a)** Fraction of postsynaptic neurons whose preferred orientation can be correctly predicted from the total excitatory input current (as in the gray line in Fig. 2), shown as a function of the number of detected inputs (in-degree). Postsynaptic neurons were divided into quartiles based on their in-degree **(b)** Fraction of cases in which the emergent orientation of a model neuron (as in Fig. 3d) matches its target orientation for different values of in-degree *K* within the estimated lower and upper bounds (see Methods).

**FIG. S2.**
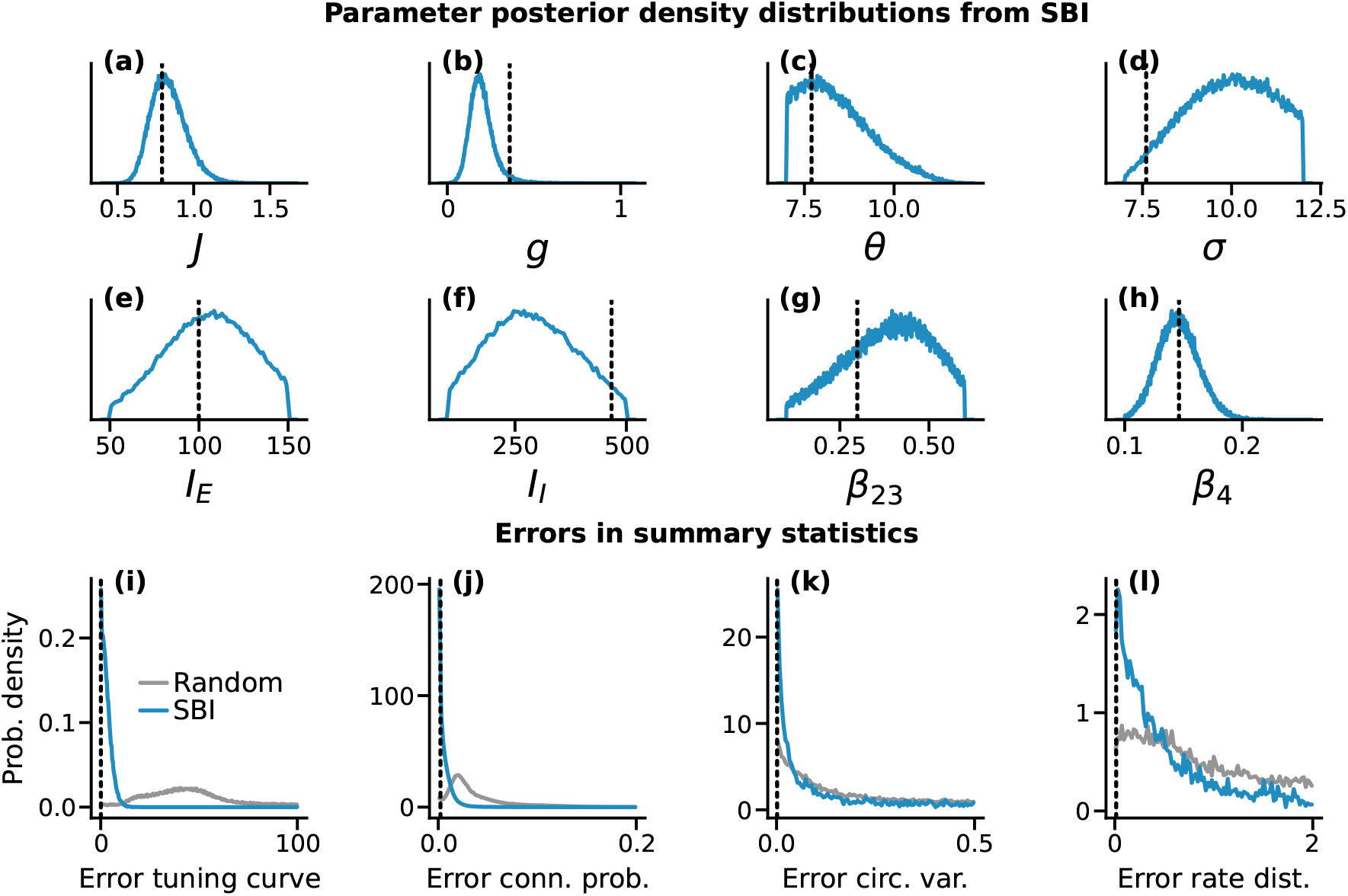
Posterior distributions of model parameters from Fig. 4. **(a–h)** Marginal posterior distributions of all fitted parameters in the connectome-constrained network model, obtained via simulation-based inference. The parameter values used for the simulations shown in Fig. 4 are indicated by vertical dashed lines. **(i–l)** Summed mean squared error (MSE) of grouped summary statistics for interpretability. **(i)** Sum of MSEs for the firing rates at 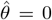 and 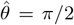. **(j)** Sum of MSEs for the decrease in connection probability at *π/*2 for layers 2/3 and 4. **(k)** Sum of MSEs for the mean, standard deviation, and skewness of the circular variance. **(l)** Sum of MSEs for the mean and standard deviation of the logarithm of firing rates.

**FIG. S3.**
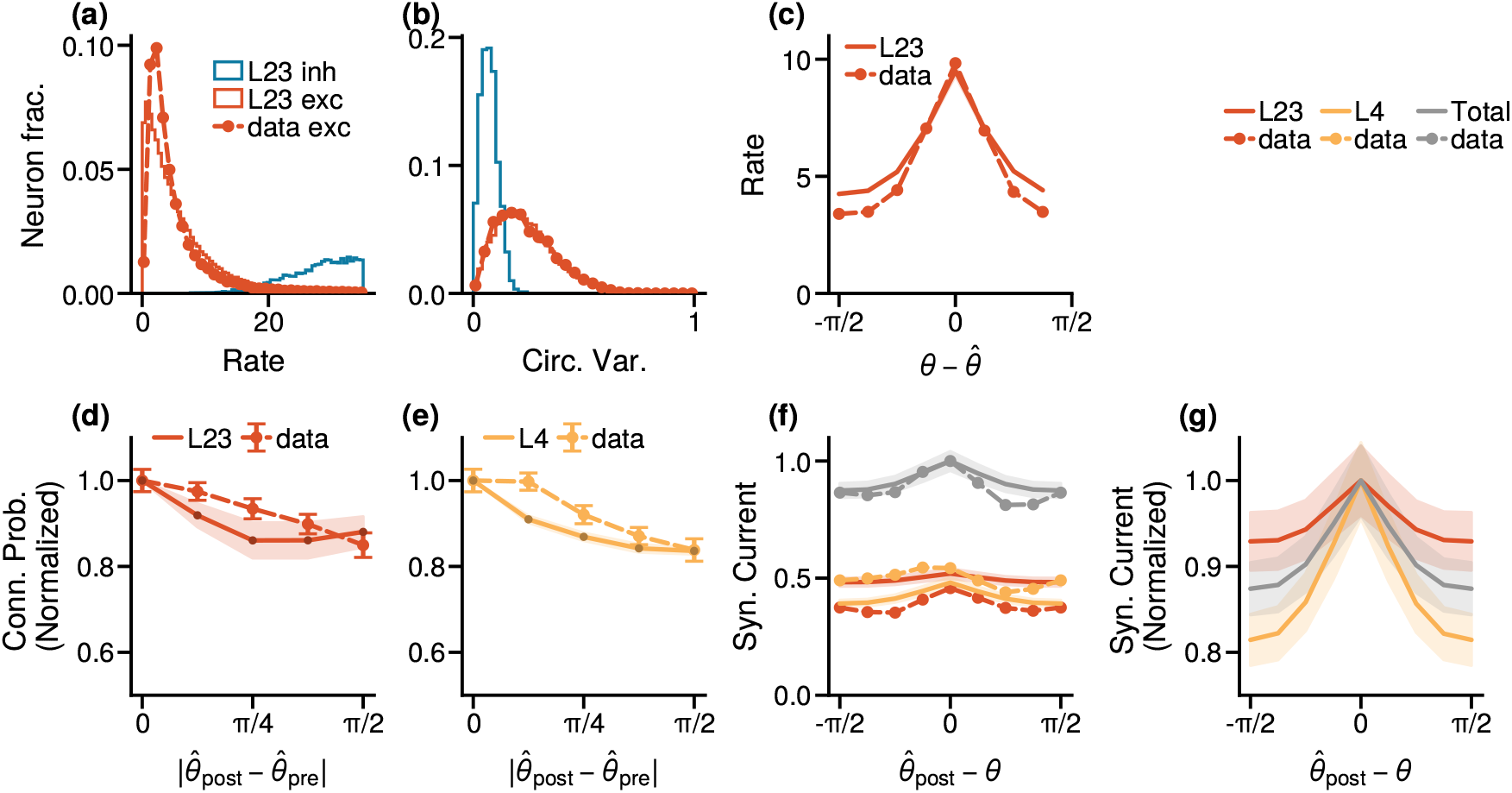
Connectome-constrained models capture diverse functional and structural features of L2/3 (with orientation-dependent inhibitory connectivity). Same analyses as in Fig. 4, but for network models in which inhibitory connections are orientation-dependent rather than random (see Methods). The main network features and selectivity measures remain qualitatively unchanged, indicating that the specific structure of inhibitory connectivity has a limited impact on overall circuit selectivity.

**FIG. S4.**
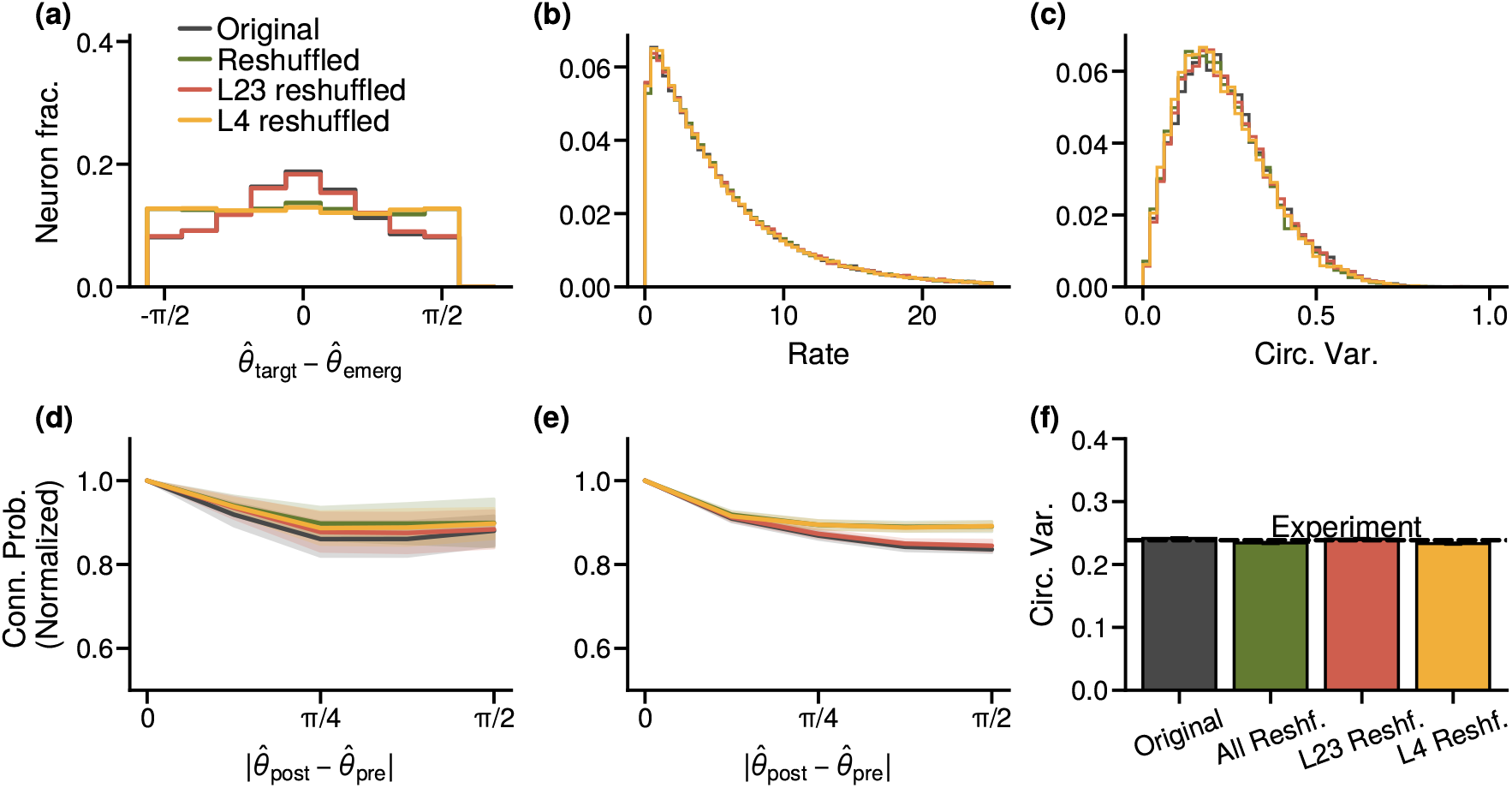
Connectivity randomization in models with orientation-dependent inhibitory connectivity. Same analyses as in Fig. 5, but for network models in which inhibitory connections depend on orientation selectivity (see Methods). Shuffling inhibitory connections produces only minor changes, similar to those observed in Fig. 5, indicating that incorporating orientation-dependent inhibitory structure does not alter the main conclusions.

**FIG. S5.**
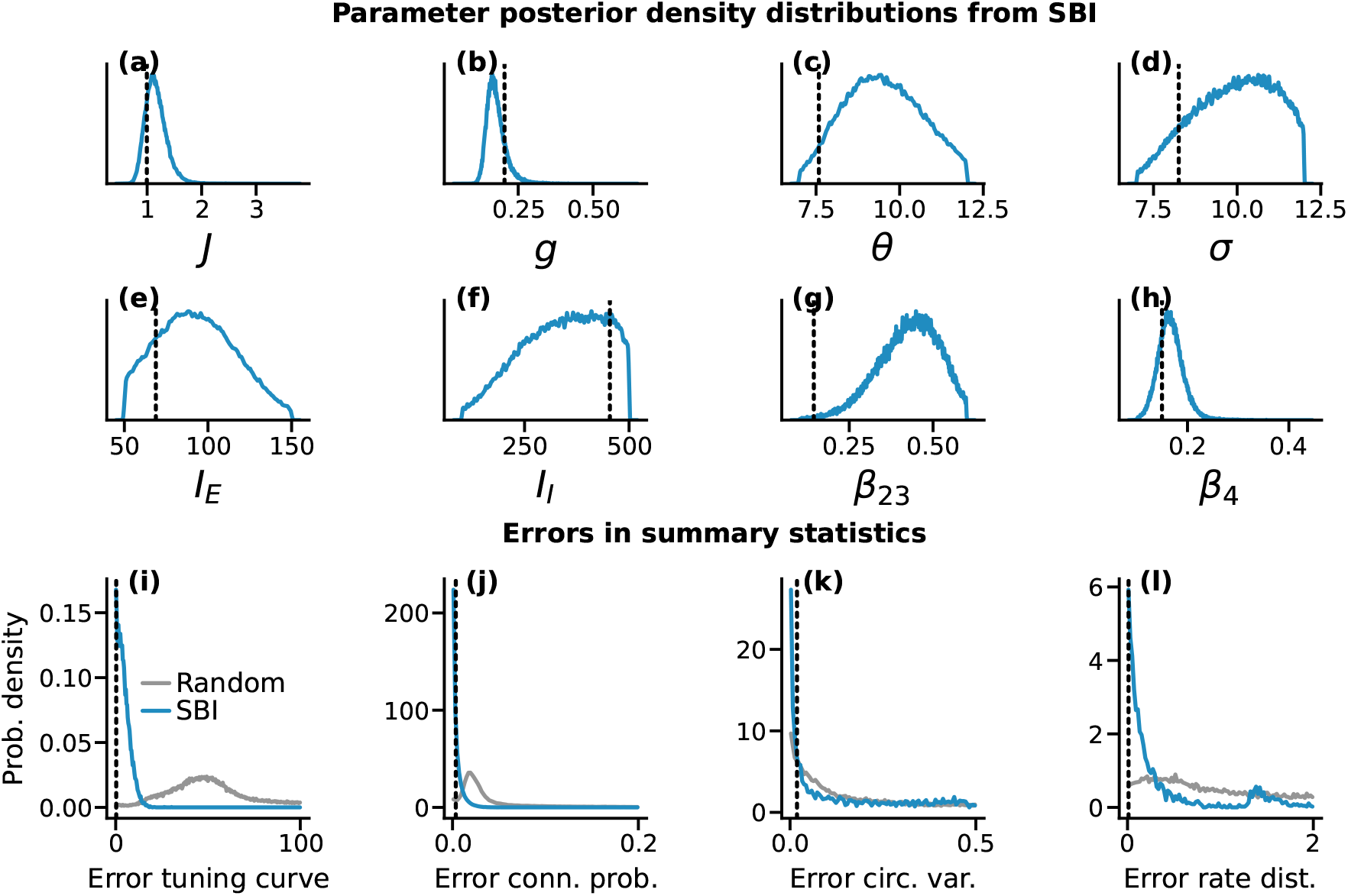
Posterior distributions for the orientation-dependent inhibitory connectivity model (related to Fig. S3). **(a-h)** Marginal posterior distributions for all fitted parameters in the model with orientation-dependent inhibitory connectivity. The values used for simulations with tuned inhibition (Sup. Fig. S3) are marked with a vertical dashed line. **(i-l)** Sum of the ean square error of selected summary statistics adequately grouped, as in Sup. Fig. S2.

**FIG. S6.**
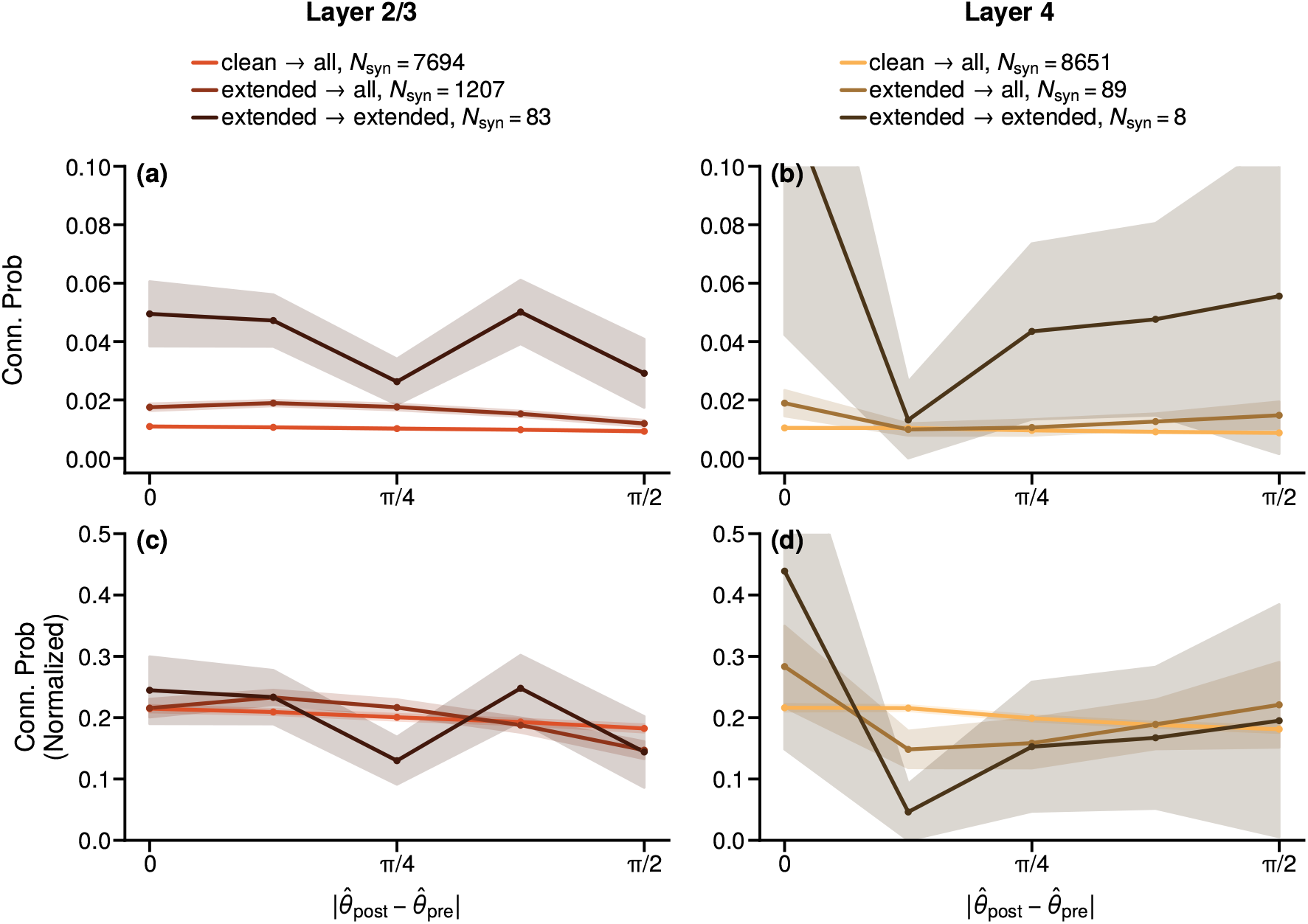
Dependence of connection probability on proofreading status. (a,b) Connection probability between pre- and postsynaptic neurons with different proofreading statuses in layer 2/3 (a) and layer 4 (b). (c,d) Same as (a,b), but each probability *p*(*θ*) is normalized to 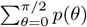 to facilitate comparison across groups. The dependence of connection probability on orientation is largely independent of proofreading status, with fluctuations mainly reflecting the limited number of available connections.

